# Chronic ethanol drinking in non-human primates induces inflammatory cathepsin gene expression in alveolar macrophages accompanied by functional defects

**DOI:** 10.1101/2021.07.30.454528

**Authors:** Sloan A. Lewis, Brianna Doratt, Suhas Sureshchandra, Allen Jankeel, Natali Newman, Kathleen A. Grant, Ilhem Messaoudi

**Author notes:** Corresponding Author: Ilhem Messaoudi, Molecular Biology and Biochemistry, University of California Irvine, 2400 Biological Sciences III, Irvine, CA 92697.

## Abstract

Chronic alcohol drinking is associated with increased susceptibility to viral and bacterial respiratory pathogens. Investigating the effects of alcohol on the lung is challenging in humans because of the complexity of human drinking behavior and the challenge of obtaining samples. In this study, we utilize a rhesus macaque model of voluntary ethanol self-administration to study the effects of alcohol on the lung in a physiologically and genetically relevant model. We report a heightened activation and inflammatory state in alveolar macrophages (AM) obtained from ethanol drinking animals that is accompanied by increased chromatin accessibility in intergenic regions that regulate inflammatory genes and contain binding motifs for transcription factors AP-1, IRF8, and NFKB p-65. In line with these transcriptional and epigenetic changes at basal state, AM from ethanol drinking animals generate elevated inflammatory mediator responses to LPS and respiratory syncytial virus (RSV). Analysis using scRNA-Seq revealed heterogeneity in lung-resident macrophage and monocyte populations, including increased abundance of activated and cathepsin-expressing clusters and accelerated differentiation with ethanol. Finally, functional assays show increased mitochondrial content in AM from ethanol drinking animals, which is associated with observed increased ROS and decreased phagocytosis capacity. This comprehensive epigenomic, transcriptional and functional profiling of lung macrophages after ethanol drinking in macaques provides previously unidentified mechanisms of ethanol induced infection susceptibility in patients with alcohol use disorders.

## INTRODUCTION

Alcohol use is prevalent in the United States with over 50% of people 18 years or older reporting alcohol consumption with the previous 30 days (National Survey on Drug Use and Health 2019). Amongst these individuals, 25% report binge drinking and 6.3% report heavy drinking. Long term heavy drinking is associated with numerous adverse health outcomes, including increased incidence of cardiac disease (1, 2), certain types of cancer (3-6), liver cirrhosis (7), and sepsis (8), making it the third leading preventable cause of death in the United States (9). Of importance, chronic heavy alcohol drinking compromises lung health and immunity leading to increased susceptibility to both bacterial and viral pulmonary infections (10), notably respiratory syncytial virus (RSV), (11) community-acquired pneumonia (12-14), and tuberculosis (15, 16). Alcohol use is also a risk factor for acute respiratory distress syndrome (ARDS) (17, 18) and can increase the risk of admission to intensive care unit (ICU) in patients with pneumonia (10, 12, 17, 19). While the mechanisms underlying increased vulnerability and severity of pulmonary infections with chronic alcohol consumption have yet to be fully elucidated, studies using rodent models as well as *in vitro* cell cultures have identified defects in the beating of the ciliated epithelium (20-22) as well as impaired epithelial barrier function (23, 24) as major risk factors. Moreover, these studies report significant defects in both the innate and adaptive branches of the immune system (10), especially within alveolar macrophages (AM), the first line of defense in the lung (25). Specifically, prolonged alcohol exposure alters the ability of AM to release cytokines and chemokines needed to recruit immune cells into the lung (26, 27) as well as their ability to clear both microbes and dying cells to reduce damage to tissue (28) potentially due to oxidative stress (29). The molecular basis for altered macrophage metabolism and function in the lung with alcohol is yet to be determined.

Lung-resident macrophages can be categorized into interstitial and alveolar with interstitial macrophages primarily residing within the tissue while alveolar macrophages are predominantly found within the lumen of the alveoli (30). Studies in mice have revealed that lung macrophages are derived from yolk sac and fetal liver as well as from bone marrow monocytes (30). It is believed that embryonically derived macrophage populations have a self-renewal capacity and are functionally distinct from the monocyte-derived macrophages populations, however, whether this is true in humans is still unanswered (30). Recent studies have uncovered enormous heterogeneity in lung macrophage populations, but many questions remain as to how environmental factors or inflammatory settings alter the functional capabilities of these cells to clear pathogens and repair tissue.

In this study, we use bronchoalveolar lavage (BAL) samples collected from rhesus macaques that voluntarily self-administered ethanol (EtOH) or an isocaloric solution for 12 months to examine the alcohol-induced epigenetic and transcriptomic changes that are coupled to altered macrophage function. We report a heightened activation state in AM obtained from EtOH drinking animals that was accompanied by increased chromatin accessibility in intergenic regions that regulate inflammatory genes and binding motifs for transcription factors AP-1, IRF8, and NFKB p-65. In line with these transcriptional and epigenetic changes at basal state, AM from EtOH drinking animals generated heightened inflammatory mediator responses to LPS and RSV. In contrast, expression of genes associated with tissue repair and antiviral type interferon responses were reduced with EtOH drinking. Additional analysis using scRNA-Seq revealed considerable heterogeneity in AM and monocyte populations, including increased abundance of activated and cathepsin-expressing cells. Finally, functional assays show reduced phagocytic capacity, but increased ROS production that may be mediated by increased mitochondrial content in AM from EtOH drinking animals. Our comprehensive epigenomic, transcriptional and functional profiling of lung macrophages after *in vivo* EtOH exposure in rhesus macaques provides novel mechanisms by which patients with alcohol use disorders have increased susceptibility to respiratory infections.

## RESULTS

### Chronic EtOH exposure alters surface activation and chemokine receptor expression on monocyte and macrophage populations in the lung

Chronic heavy alcohol drinking has been shown cause activation and hyper-inflammation in monocytes in the blood (31) and macrophages in the spleen (32). The alveolar space in the lung is home to a large population of tissue-resident macrophages and infiltrating monocytes that are the first responders to respiratory infections. Given that patients with alcohol use disorders have increased susceptibility to respiratory pathogens, we used a multipronged approach to uncover the pleiotropic impact of chronic heavy drinking on the transcriptional, epigenetic, and functional landscape of the alveolar macrophages (AM). We collected bronchial alveolar lavage (BAL) samples from male and female rhesus macaques that either consumed EtOH or an isocaloric solution for 12 months (**Figure 1A and Supp. Table 1A**). We first determined the impact of chronic EtOH on the phenotype of AM by profiling cell surface markers using flow cytometry (n=6 control, 8 EtOH). Based on previous studies (33-35), we identified alveolar macrophages (AM) as CD206+CD169+, interstitial macrophages (IM) as CD206+CD169-, and infiltrating monocytes as CD206-CD169-CD14+HLA-DR+ (**Figure 1B**). AM were further subdivided based on expression of CD163 (**Figure 1B**). No significant differences in frequencies of these major macrophage/monocyte populations were observed with EtOH exposure. (**Figure 1C**). However, examination of surface activation markers and chemokine receptors using flow cytometry showed a modest increase in CD40 and a modest decrease in CD11c expression on AM; modest increase in CCR2 expression on IM and infiltrating monocytes; and a modest increase in CD163 expression on IM with chronic EtOH consumption (p≤0.1) (**Figure 1D**). Similarly, chronic EtOH consumption led to heightened activation of the CD163lo AM subset as indicated by increased expression of CD14, HLA-DR, CD40, CD86 and CX3CR1 (p≤0.1) (**Figure 1E**). To determine whether EtOH dose impacted the expression of surface markers, we performed linear regression analyses of median fluorescence intensities (MFI) with the 12-month average dose of ethanol (g EtOH/ kg body weight/ day). Average daily EtOH drinking positively correlated with CD40, CX3CR1 and CCR7 expression on AM (**Figure 1F**) and with CCR5 and CCR2 expression on monocytes and IMs (**Figure 1F**). CD11c expression negatively correlated with EtOH dose in monocytes and AMs while CD86 expression positively correlated with EtOH dose in the CD163lo AM population (**Figure 1F**). Therefore, while EtOH drinking does not result in major subset redistribution, it impacts the activation status of lung resident macrophages and chemokine receptors associated with lung trafficking on monocytes in a dose-dependent manner.

**Figure 1:**
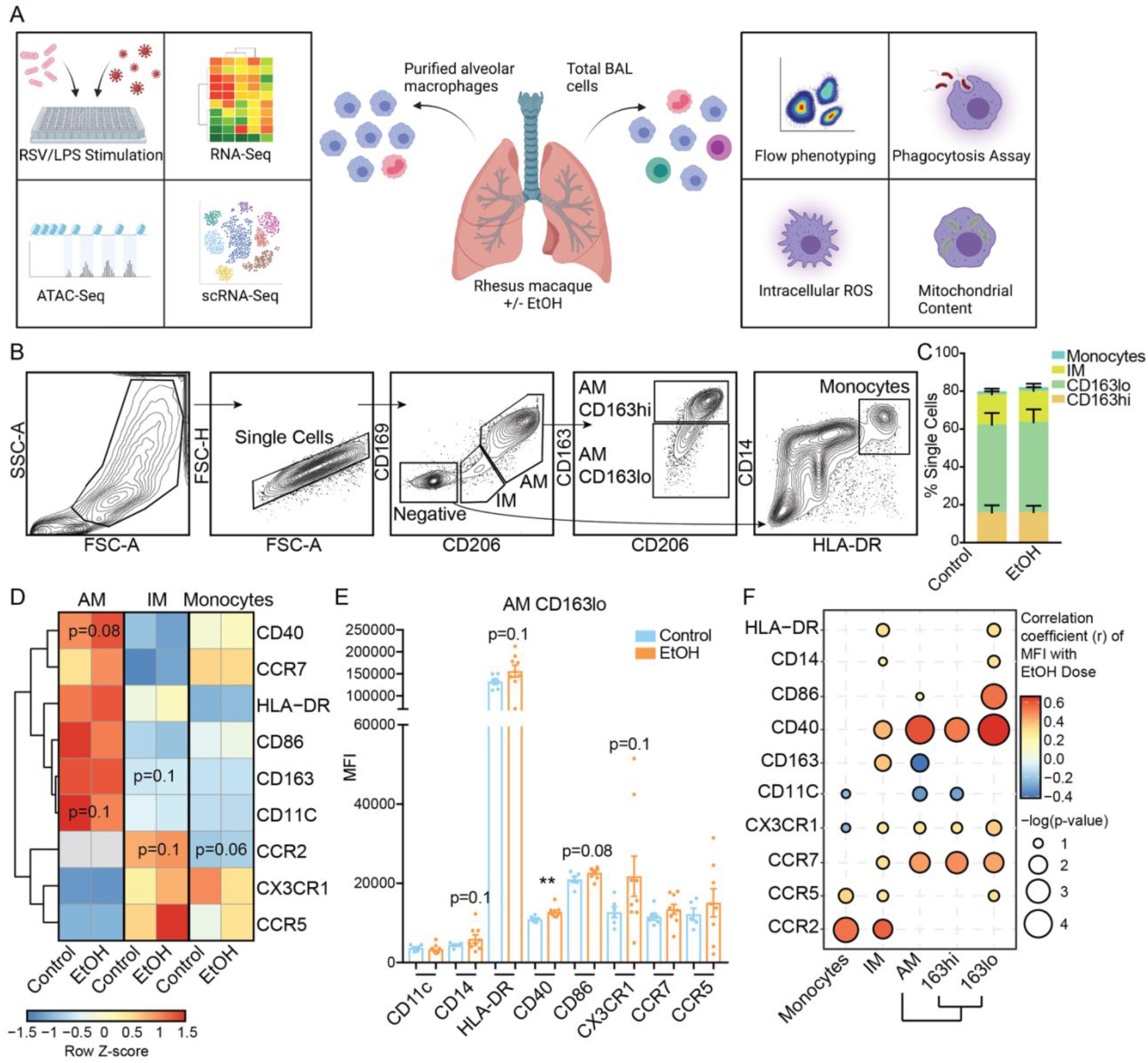
EtOH exposure alters alveolar macrophage (AM) phenotype. A) Experimental design of this study created with BioRender.com. B) Gating strategy to identify monocytes, interstitial macrophages (IM), and alveolar macrophages (AM) from bronchoalveolar lavage (BAL) samples. C) Relative distributions of the three myeloid cell subsets in the BAL. D) Heatmap showing averaged median fluorescence intensity (MFI) values for cell surface markers in each indicated subset/group. The scale is row Z-score where blue is lower and red higher expression. E) Median fluorescence intensity (MFI) of activation and chemokine surface markers measured from the CD163lo AM population. F) Bubble plot representing correlations between cell surface markers and ethanol dose in the indicated cell populations. The size of each circle represents the indicates the -log_10_ transformed p-value significance measured by linear regression. The color denotes the correlation coefficient (r) calculated by linear regression. Significance for two-group comparison was calculated by t-test with Welch’s correction where trending values are shown.

### Chronic EtOH is associated with downregulation of genes involved in tissue maintenance and wound healing in AM

We have previously reported a disruption of transcriptional programs of both peripheral monocytes and splenic macrophages with chronic alcohol consumption (31, 36). Therefore, we next examined transcriptional rewiring of lung-resident alveolar macrophages, which harbor a significant proportion of embryonically derived, self-renewing tissue-resident cells (30). AM were purified from control and EtOH animals (n=3/group) and bulk RNA sequencing performed (**Figure 1A**). EtOH exposure explained the most variability in baseline transcriptional profiles in AMs (**Figure 2A**). Differential analysis revealed 24 genes to be upregulated with EtOH (**Figure 2B**) including *CTSG, SNAP25*, and *HEBP2* which are all associated with granulocyte activation and degranulation (**Figure 2C**). EtOH consumption was also associated with 195 downregulated DEG, among which *CLEC1B*, which is associated with an anti-inflammatory macrophage phenotype, was the most significant (**Figure 2A**) (37). The downregulated DEG mapped significantly to response to wounding (*CLEC1B, NRP1, PTK2*, and *PRKACB*), cell morphogenesis (*MYO7A, PLCGG1*), and vasculature development (*HMGA2, ACTA2*) pathways (**Figure 2D and 2E**). These observations indicate that EtOH consumption skews AM away from tissue maintenance and repair and towards inflammatory responses.

**Figure 2:**
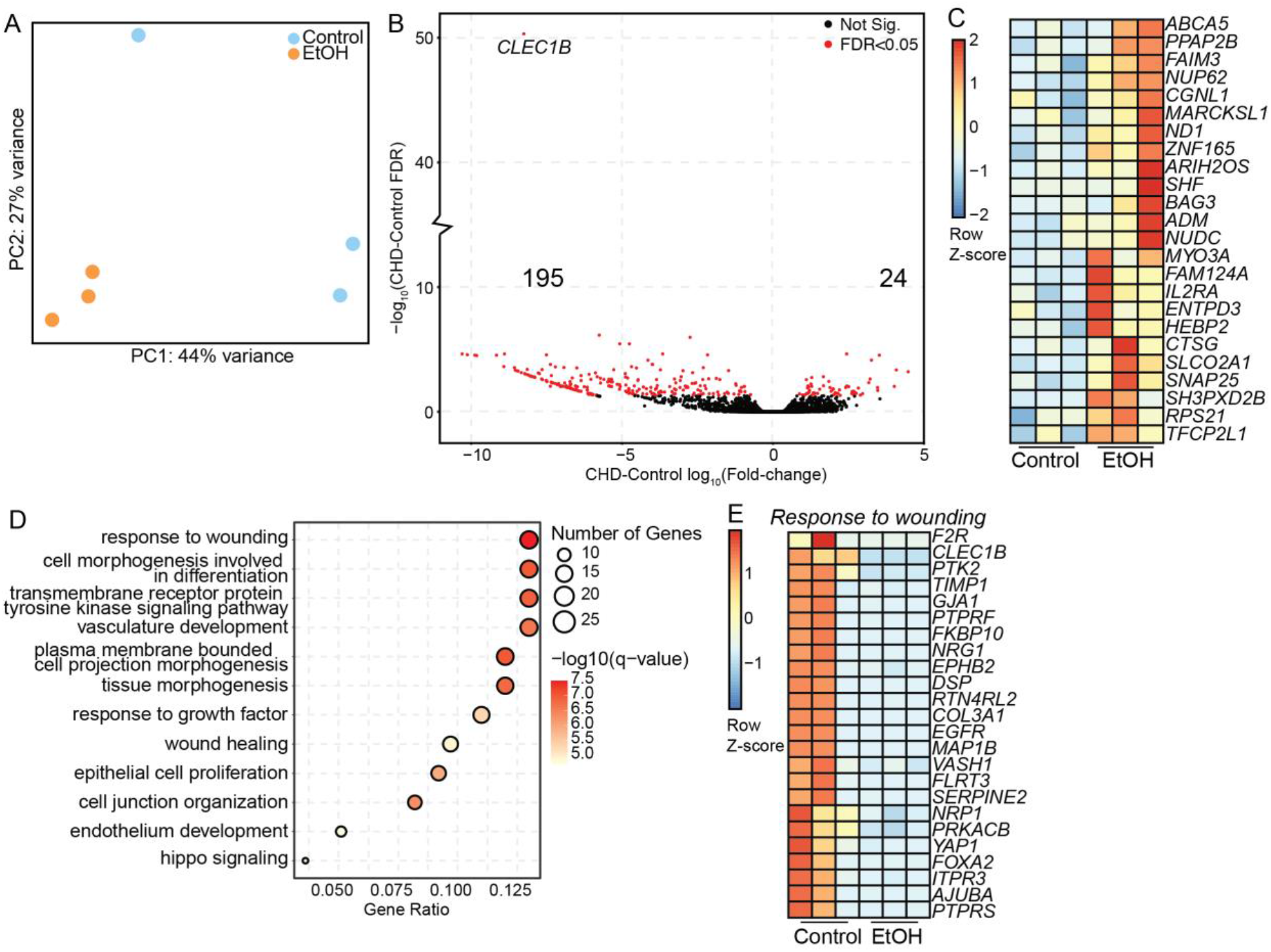
EtOH exposure induces changes in the alveolar macrophage (AM) transcriptome. CD206+ alveolar macrophages (n=3/group) were purified from total BAL using FACS and subjected to bulk RNA-Seq. A) Principal component analysis (PCA) of bulk RNA-Seq libraries from controls and EtOH AM. B) Volcano plot representing the up- and downregulated differentially expressed genes (FDR<0.05) where the X-axis is log_10_ fold-change and the Y-axis is -log_10_ FDR. C) Heatmap representing the expression of DEG upregulated with EtOH where the scale is Row Z-score representing low (blue) and high (red) expression. D) Bubble plot showing GO Biological Process enrichment of downregulated DEG with EtOH. The size of the bubble represents the number of genes associated with that term, the color represents -log_10_ q-value, and the X-axis is the ratio of genes mapping to that term to total genes. E) Heatmap representing the expression of DEG from response to wounding term where the scale is Row Z-score representing low (blue) and high (red) expression.

### EtOH exposure results in opened promoter regions at CTSG and SNAP25 genes involved in degranulation and inflammation in alveolar macrophages (AM)

To assess whether baseline transcriptional changes in the AM could be due to epigenetic changes caused by EtOH exposure, we performed ATAC-Seq on purified AM from control and ETOH animals (n=3/group). Although the relative distribution of open promoter and distal regions was comparable between controls and EtOH AM (**Figure 3A**), several differentially accessible regions were identified within the promoter and distal intergenic regions (**Figure 3B**). The 70 genes associated with promoters that were more closed with EtOH enriched to gene ontology (GO) terms associated with barrier function such as endothelium development (*CLDN3, CLDN5*) and T-helper cell differentiation (*FOXP1*) (**Figure 3C**). The 25 genes associated with promoters that were more accessible with EtOH mapped to GO term regulation of hormone levels (*CTSG, SNAP25*, and *MYO3A*) (**Figure 3C, D**). Intriguingly, these 3 genes were also upregulated based on the bulk RNA-Seq analysis (**Figure 3D**).

**Figure 3:**
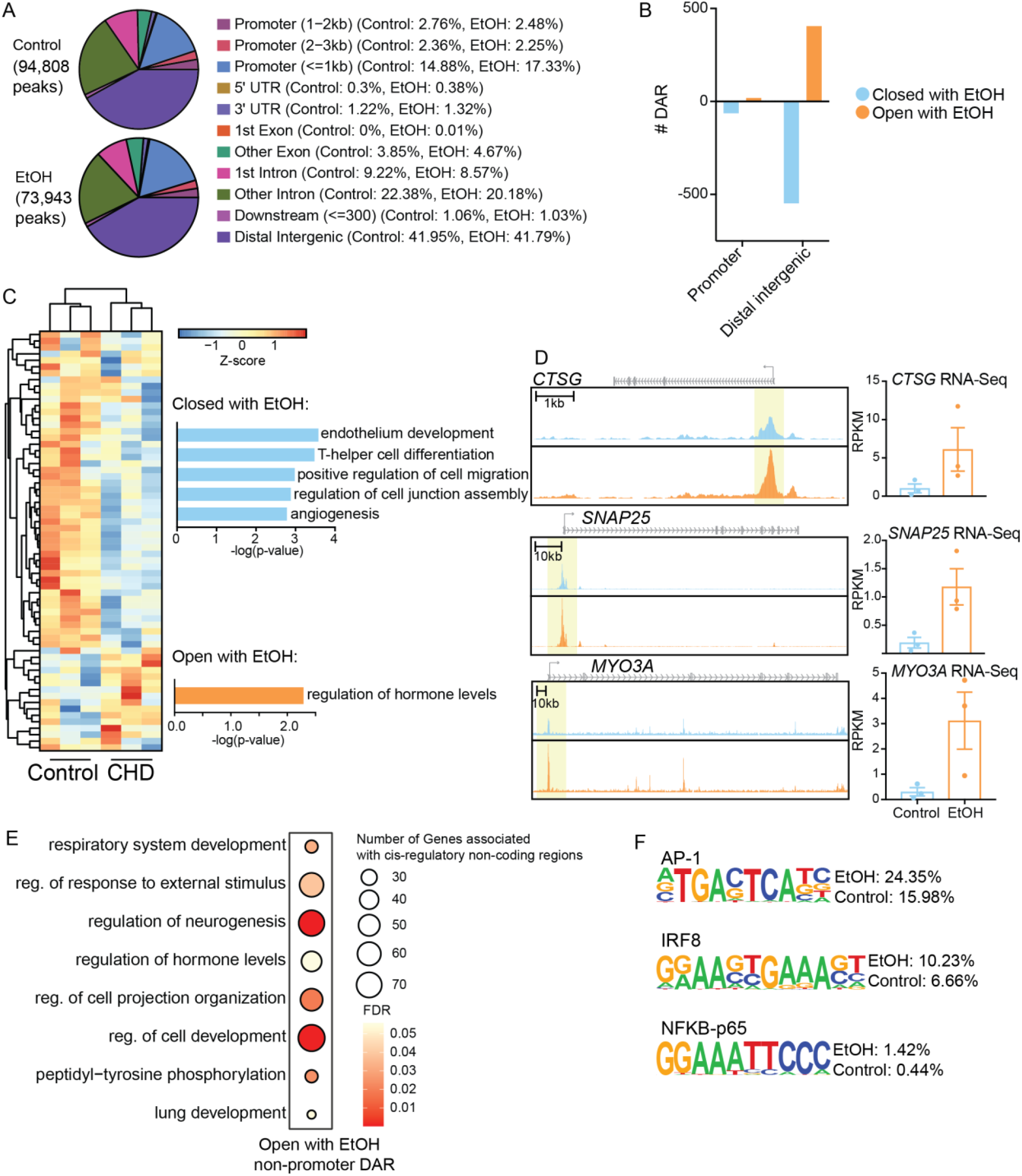
Epigenomic analysis of alveolar macrophages (AM) with chronic EtOH exposure. Alveolar macrophages (n=3/group) were purified from total BAL by CD206+ sort and nuclei were subjected to ATAC-Seq. A) Pie charts showing genomic feature distribution of the open chromatin regions (fold-change ≥ 2, FDR ≤ 0.05) in control and EtOH AM. B) Bar chart showing the number of open and closed differentially accessible regions (DAR) (FDR ≤ 0.01) with EtOH in promoter and distal intergenic regions. C) Heatmap representing the open and closed differentially accessible promoter region counts where the scale is Row Z-score representing low (blue) and high (red) expression. Bar plots to the right represent the functional enrichment of those promoter DAR where the X-axis is -log_10_ p-value. D) Pile-ups of selected promoter DAR more open with EtOH. Scale is indicated. To the right of each is a bar chart of the RPKM expression value of the gene from bulk RNA-Seq analysis. E) Non-promoter DAR were lifted over to the human genome and enriched for cis-regulatory mechanisms using GREAT. Bubble plot of the open non-promoter DAR enrichment where the size of the bubble represents the number of gene regions associated with that term and the color represents the FDR significance. F) Homer motif enrichment of the distal intergenic DAR. All listed motifs have significantly enriched binding sites in the open and closed non-promoter DAR where the percentage value listed is the percentage of target sequences with that motif.

Analysis of potential cis-regulatory mechanisms of regulation in the non-promoter regions was performed by first lifting the genomic regions from the macaque to human genomes followed by enrichment using the GREAT database. This analysis revealed no significant enrichment of the regions that were less accessible with chronic EtOH; however, intergenic regions that were more accessible with chronic EtOH significantly enriched to respiratory system development (*CTGF, EGFR, TGFBR2*) and regulation of response to external stimulus (*C1QB, CD180, CXCR4, IL21*) (**Figure 3E**). Finally, we performed transcription factor (TF) binding motif analysis on the distal intergenic DAR, which showed higher likelihood of binding sites for TF that play a critical role in inflammation, notably AP-1, IRF8, and NFKB p-65 with chronic EtOH (**Figure 3F**). These observations indicate significant remodeling of the epigenetic and transcriptional landscape of AM towards a heightened inflammatory state with chronic EtOH drinking.

### AM functional response to pathogens is characterized by non-specific inflammatory mediator production, but compromised interferon transcriptional response with chronic EtOH

Previous studies have reported an exaggerated inflammatory response by myeloid cells to LPS (31, 32, 36, 38). Thus, FACS purified AM were stimulated with LPS (n=6/group), and immune mediator production was determined by Luminex (**Supp. Figure 1A and Supp. Table 2**). As described for peripheral blood and splenic macrophages, AM from EtOH drinking animals mounted a hyper-inflammatory response as indicated by heightened production of cytokines (IL-6, TNFα) and chemokines (CXCL8, CXCL10, CCL2, CCL4) compared to control AM (**Supp. Figure 1A**). We next examined responses of AM to a respiratory pathogen. To that end, FACS purified AM were stimulated with respiratory syncytial virus (RSV) *ex vivo* (n=8/group) and antiviral responses were determined using RNA-Seq and Luminex. We found broadly that in response to RSV stimulation, AM produced a majority growth factors (e.g. BDNF, VEGF, PDGF, and FGF), a few chemokines (CCL5 and CXCL10) and canonical inflammatory marker IL-6 (**Figure 4A**). Despite comparable viral loads (**Supp. Figure 1B**), AM from EtOH exposed animals also produced significantly increased amounts of additional inflammatory mediators (IL-1β, IL-12, IL-15, IFNβ) as well as other cytokines and growth factors (GM-CSF, C-CSF, IL-7) relative to their unstimulated condition (**Figure 4A**). AM from EtOH exposed animals also produced significantly higher levels of IL-6, IL-12 and TGFα relative to their control counterparts (**Figure 4B**). Moreover, significant positive correlations between IL-6, IL-12, and CCL5 concentration and EtOH dose was observed (**Figure 4C**).

**Figure 4:**
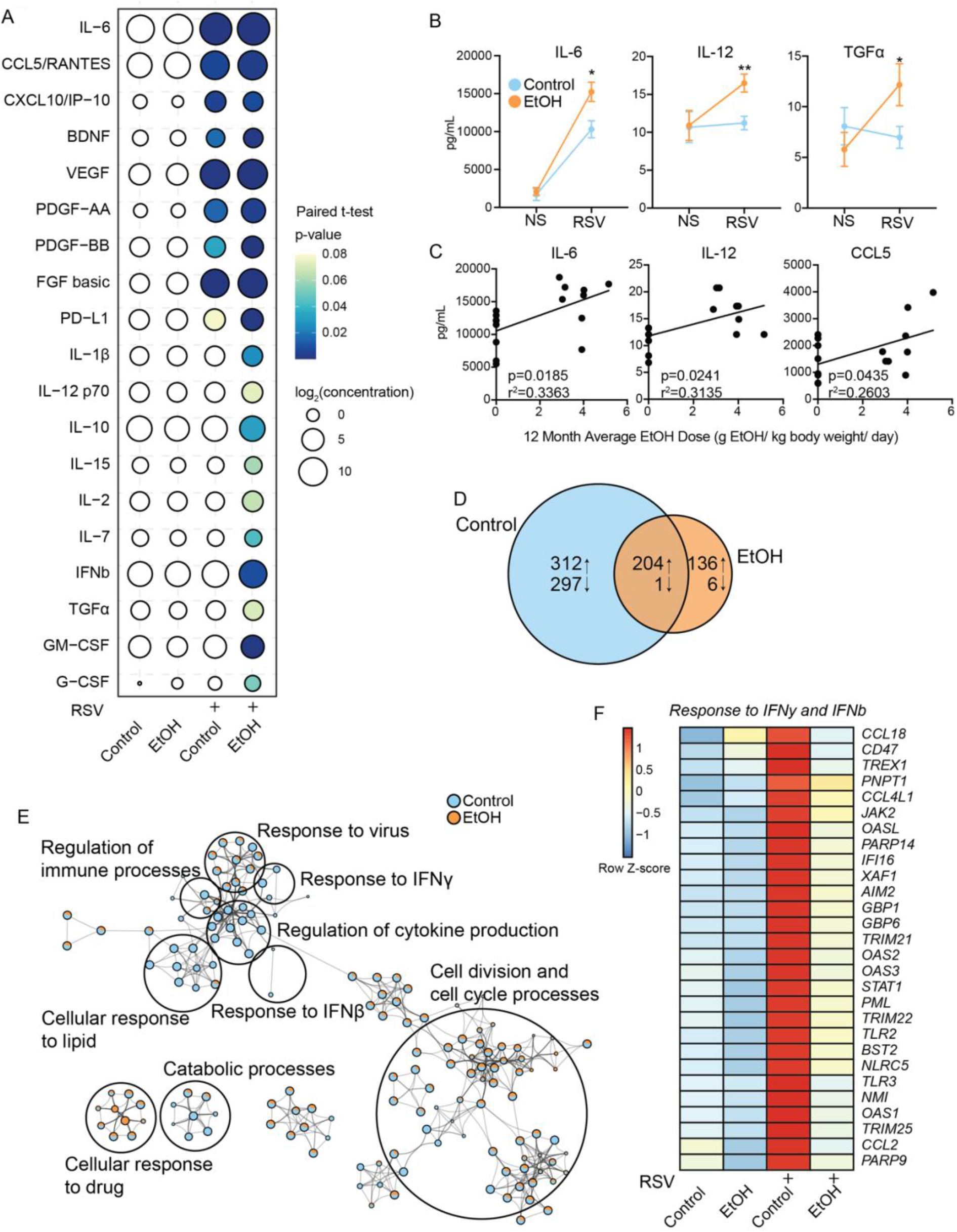
EtOH-induced non-specific inflammatory response to RSV. AM were purified and stimulated with RSV for 16 hours followed by Luminex analysis of mediator production and RNA-Seq. A) Bubble plot representing immune factor production (pg/ml) in the presence or absence of respiratory syncytial virus (RSV) by AM from control and EtOH animals. The size of each circle represents the indicates the log_2_ average concentration of the indicated secreted factor and the color denotes the p-value significance with the darkest blue representing the most significant value. The p-values were calculated between the unstimulated and stimulated conditions for each group using paired t-test. White circles indicate uncalculated or non-significant p-value. B) Line plots representing the Luminex data for the selected analytes. Significance between groups was tested by one-way ANOVA. C) Scatter plots showing linear regression analysis of selected factor concentration with dose of ethanol (g EtOH/ kg body weight/ day). P-value and r^2^ values are indicated. D) Venn diagram comparing up- and downregulated DEG after RSV stimulation in control and EtOH AM. E) Cytoscape plot of functional enrichment to GO Biological Processes of upregulated DEG from both groups. The size of the dot represen ts the number of genes enriching to that term and the pie chart filling represents the contribution of DEG from each group. Related processes are group into the larger terms circled. F) Heatmap representing the averaged expression of DEG in each group/stimulation condition from the *Response to IFNy* and *Response to IFNb* GO terms where the scale is Row Z-score representing low (blue) and high (red) expression. *=p<0.05, **=p<0.01, ***=p<0.001, ****=p<0.0001.

In contrast to the immune mediator production profile, AM from the EtOH group had a smaller transcriptional response and fewer DEG (516 DEG in control vs. 340 DEG in EtOH group) than controls (**Figure 4D**). DEG upregulated in the control group enriched to in anti-viral signaling and regulation of cytokine production processes (**Figure 4E**), whereas those upregulated in the EtOH group enriched to cell cycle and response to drug processes (**Figure 4E and Supp. Figure 1C**). Interestingly, gene signatures of cellular response to type I (IFNβ) and type II (IFNγ) interferons (*CCL18, IFI16, TLR2*, and *TLR3*) **(Figure 4F)** and immune activation (*CD80, CD86*, and *CCL2*) were upregulated only in control AM (**Supp. Figure 1D**). A significant number (297) of downregulated DEG from the control AM mapped to regulated exocytosis, cell morphogenesis, and response to wounding pathways (**Supp. Figure 1E**) such as *SIGLEC10, PTPN6*, and *SDC1* (**Supp. Figure 1F**). To determine regulatory mechanisms for the differences in transcriptional response to RSV, we used the Chea3 database (39) to predict transcription factor (TF) regulation. This analysis showed that genes upregulated only in control AM with RSV were regulated by phagocytosis and viral response associated TFs PLSCR1, SP100, and IRF7, while genes upregulated in EtOH AM were regulated by inflammatory TF HMGA2 (**Supp. Figure 1G**). These observations indicate that chronic EtOH drinking results in non-specific inflammation coupled with dysfunctional anti-microbial responses in AM.

### scRNA-Seq profiling reveals significant changes in alveolar macrophage (AM) cell states with chronic EtOH

To investigate the impact of chronic EtOH on AM cell states, we performed scRNA-Seq on CD14+ purified cells from BAL samples obtained from control and EtOH animals (n=3/group) (**Supp. Figure 2A**). Uniform manifold projection (UMAP) of clustering analysis revealed 10 clusters (**Figure 5A,B**). To identify infiltrating blood-derived monocytes, we integrated single cell profiles of BAL macrophages with those of blood monocytes from the same animals (31) (**Supp. Figure 2B**). We projected cells that clustered with the blood monocytes back onto the UMAP, which revealed that cluster 4 was the major monocyte subset with some monocyte infiltration in cluster 7 (**Supp. Figure 2C**). Expression of major macrophage/monocyte markers grouped cells into tissue resident macrophages (TRM 0, 1, 2, 8; expressing high levels of *FABP4, CD163*, and *MRC1, SIGLEC1*), monocytes (4; expressing high levels of *CD14, IL1B*, and *CCL2*), and monocyte-derived macrophages (MDM 3, 5, 6, 7, 9; intermediate expression of TRM and monocyte markers) (**Supp. Figure 2D**). TRM could be further divided into four clusters based on expression of *CYBB* (cluster 0), *S100A10* (cluster 1), *CD48* (cluster 2), and *MKI67* (cluster 8, proliferating) (**Figure 5C and Supp. Table 3**). MDM were divided into five clusters based on expression of *CFD* (cluster 3), *CTSD* (cluster 5, cathepsin high), *MSMO1*(cluster 6), *ISG15* (cluster 7 viral, monocyte infiltration), and *CXCL1* (cluster 9, activated) (**Figure 5C**). Cells from EtOH animals almost exclusively made up the activated MDM cluster 9 and were more enriched in TRM cluster 1 and MDM cluster 5 (cathepsin high) (**Figure 5D**). On the other hand, cells from controls were more abundant in the MDM cluster 3 and blood monocyte cluster 4 (**Figure 5D**).

**Figure 5:**
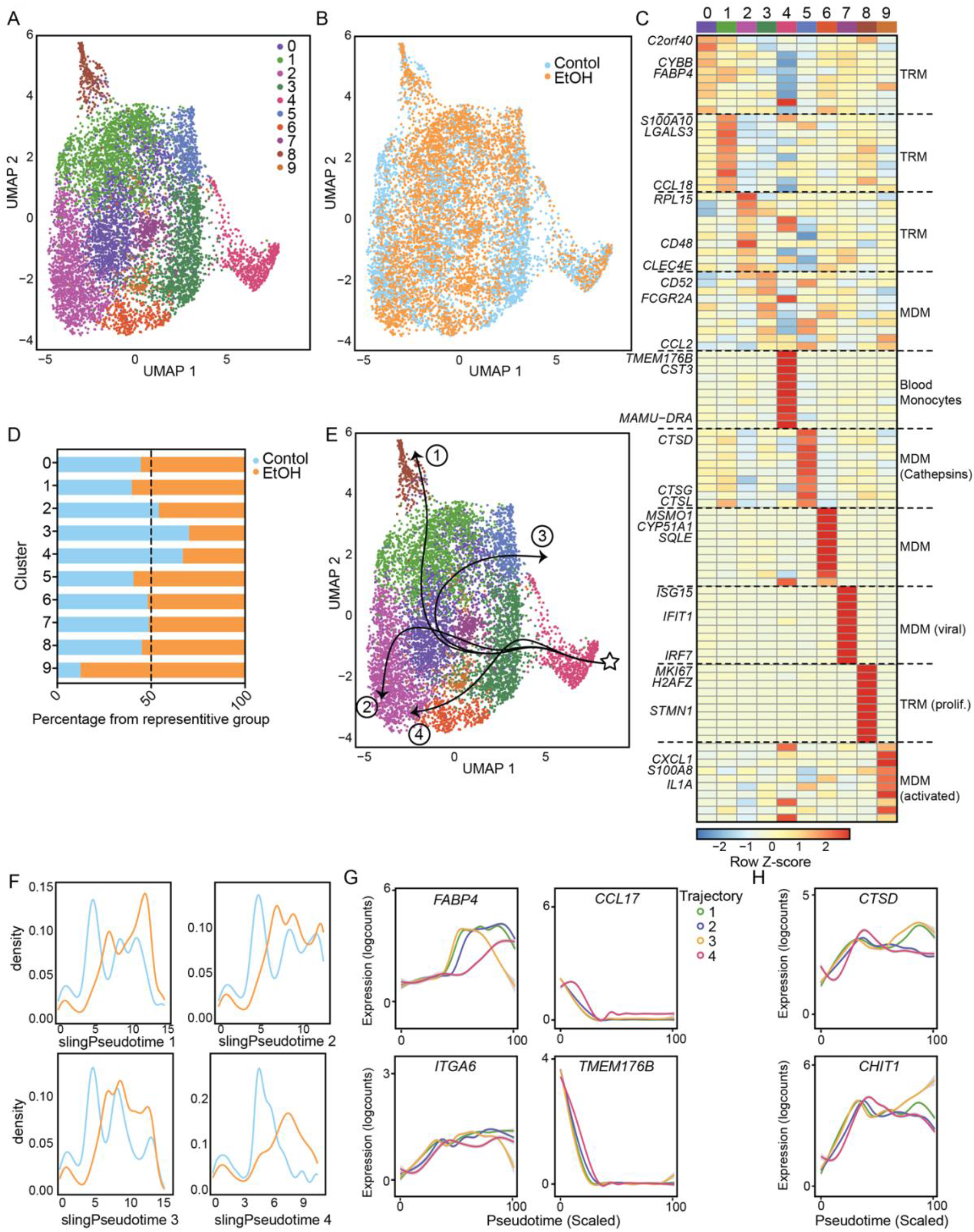
scRNA-Seq profiling of alveolar macrophages after EtOH exposure. Macrophages and monocytes (n=3 control/ 3 EtOH) were purified from total BAL and subjected to 10X scRNA-Seq analysis. A,B) Visualization of total cells by uniform manifold approximation and projection (UMAP) colored by cluster (A) and by group (B). C) Heatmap representing averaged expression of cluster marker genes identified using the *FindMarkers* function where the scale is Row Z-score representing low (blue) and high (red) expression. D) Relative distribution of the cells from control (blue) or EtOH (orange) groups within each identified cluster. E) UMAP with indicated pseudotime lineages identified by Slingshot trajectory analysis. F) Cell density plots for Control and EtOH groups across each of four trajectory lineages determined by Slingshot. G,H) Log expression of FABP4, ITGA6, CCL17, TMEM176B (G), CTSD, and CHIT1 (H) plotted for each cell across the indicated scaled slingshot pseudotime trajectory (trendline shown).

Next, we performed trajectory analysis using Slingshot and identified 4 unique trajectory paths starting from blood monocytes the culminated into clusters 7, 2, 5, and 6 (**Figure 5E**). Cells from control animals were more abundant at the start of the trajectory while cells from EtOH animals were more abundant at the end suggesting accelerated differentiation of monocytes with chronic EtOH consumption (**Figure 5F**). All trajectories were characterized by increased expression of *FABP4* and *ITGA6* and decreased expression of *CCL17* and *TMEM176B* indicative of the transition from blood to tissue resident cells (**Figure 5G**). Interestingly, trajectory 3 had decreased expression of *FABP4* and *ITGA6* but increased expression of *CTSD* and *CHIT1* at the end of the pseudotime suggesting a heightened inflammatory state (**Figure 5G,H**). Additional differential analyses on the major TRM subsets (0, 1, 2) showed increased expression of inflammatory *CCL2* but decreased expression of *FABP4* with EtOH in these clusters (**Supp. Figure 2E**). We conclude that EtOH induces subset redistribution within the AM, accelerating the differentiation from monocytes to macrophage subsets with high expression of inflammatory cathepsins.

### EtOH-induced increase in mitochondria skews alveolar macrophages towards hypoxia and ROS production and away from phagocytic and antigen presentation processes

To assess the functional implications of EtOH-induced changes in cell states, we carried out functional enrichment of the gene markers of the clusters that were more abundant in EtOH animals (TRM cluster 1, MDM cluster 5, MDM cluster 9). We identified that all three clusters mapped significantly to neutrophil degranulation, myeloid leukocyte activation, and regulated exocytosis (**Figure 6A**). All marker genes enriched to GO terms associated with myeloid cell activation, response to drug, and exocytosis (**Figure 6A**). This enrichment was most evident for MDM cluster 9 marker genes (**Figure 6A**). Additionally, TRM1 and MDM9 cluster gene markers mapped to response to hypoxia and positive regulation of cell migration (**Figure 6A**). To follow up on these observations, we performed module scoring. We found phagocytosis, antigen presentation, and cell adhesion to be downregulated with EtOH whereas HIF1A signaling, chemokine signaling, and cytokine signaling were upregulated (**Figure 6B,C and Supp. Table 4**).

**Figure 6:**
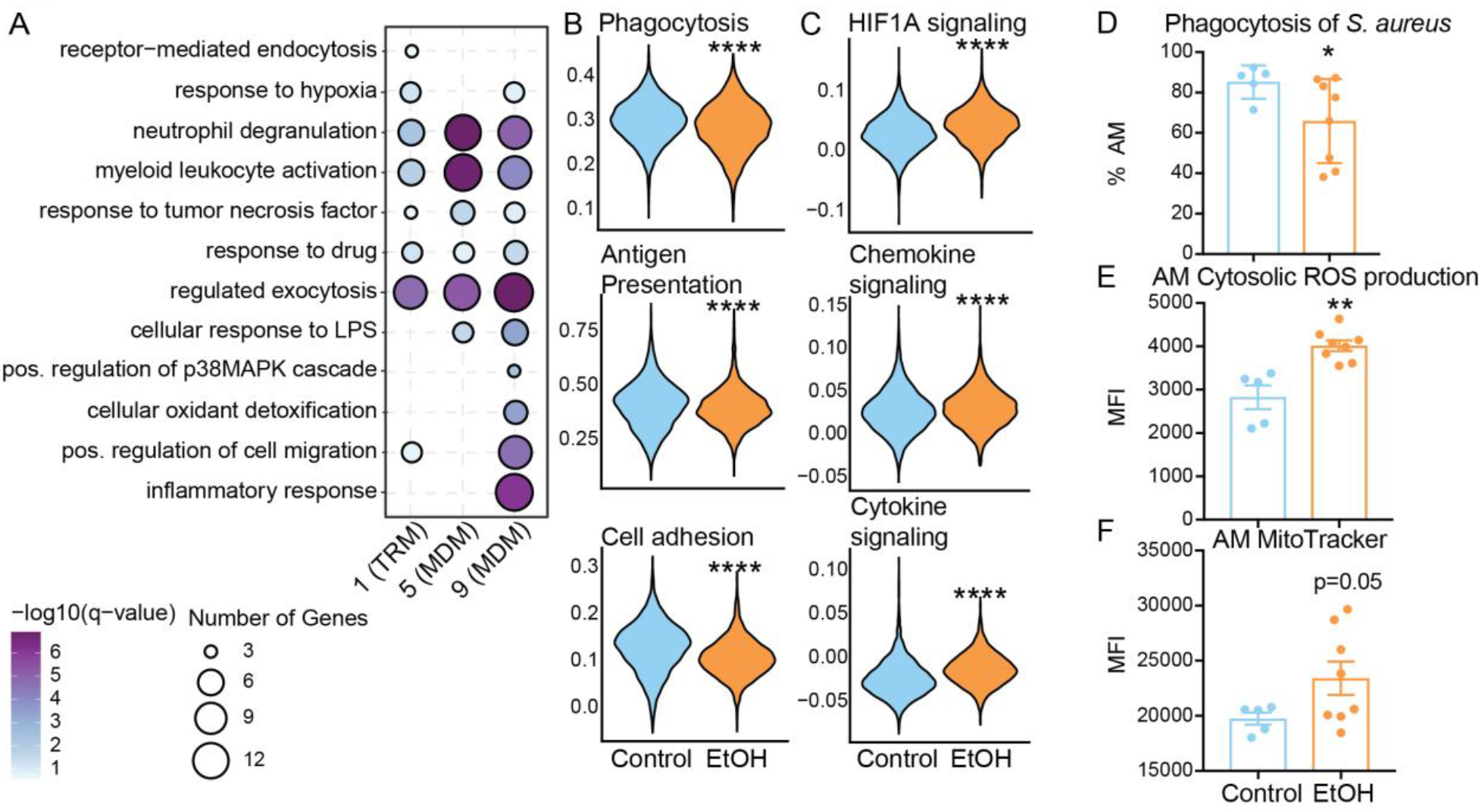
Functional implications of EtOH exposure on alveolar macrophages (AM) A) Bubble plot showing functional enrichment of genes highly expressed in clusters 1, 5, and 9. The size of the bubble represents the number of genes in that term and the color represents the -log_10_ q-value significance. B,C) Violin plots representing module scoring of total cells from each group. Significance was calculated using Mann-Whitney test. D) Bar plot representing percentage of AM positive for pHrodo Red *S. aureus* particles. E) Bar plot showing median fluorescence intensity (MFI) of intracellular oxidative stress stained by CellROX Green Reagent in AM. F) Bar plot showing median fluorescence intensity (MFI) of intracellular mitochondria stained with MitoTracker Red in AM. *=p<0.05, **=p<0.01, ***=p<0.001, ****=p<0.0001.

To further define the impact of these transcriptional changes, we incubated BAL cells with *S. aureus* labeled pHrodo and measured phagocytic capacity of AM using flow cytometry. AM from EtOH exposed animals exhibited a reduced capacity for phagocytosing bacteria compared to controls (**Figure 6D**). Additionally, cytosolic ROS production was increased in AM and IM after LPS stimulation with EtOH indicating a heightened oxidative state (**Figure 6E and Supp. Figure 2F**). Finally, as ROS production and activation in macrophages have been linked to mitochondrial function (40), we profiled the intracellular mitochondria content in AM and IM and identified significantly increased mitochondria with EtOH in AM (**Figure 6F and Supp. Figure 2G**). Given heightened inflammatory responses in the AM, we sought to determine whether alcohol drinking affected the M1/M2 polarization of the lung myeloid cells. We performed module scoring of M1 and M2 genes (41) and identified that EtOH significantly skews the monocytes and macrophages towards an M1-like and away from an M2-like phenotype (**Supp. Figure 2H**). Altogether, these data indicate that increased levels of mitochondria in AM with EtOH may be skewing the AM towards hypoxia and ROS production and away from phagocytic functional processes.

## DISCUSSION

Tissue resident macrophage and infiltrating monocyte populations make up a majority of the immune cells in the alveolar space where they interact with insults to the respiratory tract including toxins, pathogens and allergens (42). They are responsible for mounting an inflammatory immune response when necessary and moreover remodeling and repairing the tissue. Under homeostatic conditions, a tight balance between inflammatory and anti-inflammatory responses is maintained. This delicate balance can be dysregulated by environmental factors including pollutants, smoking, and alcohol drinking (43). Indeed, alcohol consumption results in increased susceptibility to respiratory diseases, but the mechanism underlying this increased vulnerability are not completely understood. Therefore, in this study, we carried out a comprehensive examination of the impact of chronic heavy alcohol consumption on the transcriptome, epigenome, and function of AM obtained from a rhesus model of voluntary ethanol self-administration. Specifically, bronchoalveolar lavages (BAL) were obtained after 12 months of drinking. This process is known to capture AM and infiltrating monocyte populations but not interstitial macrophages (IM) that reside within the lung tissue. However, our flow cytometry data indicate the presence of a small fraction of IM (CD206+CD169) in the BAL samples in addition to a large AM population. Broadly, our findings show that ethanol drinking leads to a heightened activation and inflammatory state in alveolar macrophages accompanied by reduced functional abilities.

A prominent observation of this study was increased expression of cathepsin G (*CTSG*) in AM that was accompanied by increased chromatin accessibility at *CTSG* promoter with chronic EtOH drinking. Furthermore, scRNA-Seq analysis revealed the presence of a cluster of monocyte derived macrophages dominated by cells from the EtOH group with heightened expression of cathepsins *CTSD, CTSG*, and *CTSL*. Cathepsins are proteases that are active in low pH lysosomes and have versatile functions in innate immunity, activation, tissue degradation (44). Dysregulated expression of cathepsins has been linked to diseases including arthritis, muscular dystrophy, and tuberculosis (44). The significance of increased cathepsin expression by ethanol in AM needs to be further studied to determine its importance in AM function and lung immunity. It has been previously reported that AM from patients with alcohol use disorders have elevated inflammatory mediator expression (45, 46). To complement this, we identified chronic EtOH drinking in macaques resulted in increased activation surface marker expression on AM as indicated by significantly increased CD40 on CD163lo AM. As CD163 is associated with an M2 or resolving macrophage phenotype (47), this indicates skewing towards a more inflammatory macrophage population. We additionally noted positive correlations of chemokine receptors CCR7, and CX3CR1 with EtOH dose in the AM populations. CCR7 has been associated with an M1-like phenotype in AM (48), and CX3CR1 has been implicated in TNFα and IL-6 production in tissue resident mononuclear phagocytes in the lungs of smokers (49) as well as in profibrotic macrophage subsets (50). This skewing towards M1-like phenotype was confirmed by scRNA-Seq data where the module score of genes associated with an M1-like phenotype was higher while that of M2-like phenotype was lower in the ethanol group. Additionally, transcription factor motif analysis revealed enrichment of binding sites for pro-inflammatory TF AP-1, IRF8, and NFκB with ethanol. These data are in line with our previously reported changes in splenic macrophages and circulating monocytes indicating broad epigenetic rewiring by *in vivo* chronic drinking (31, 32). Collectively, these cell surface and epigenetic alterations at baseline state in AM from ethanol drinking macaques could lead to heightened inflammation in the lung environment, which could further lead to tissue damage and risk of infection.

Previous studies have identified altered production of cytokines and chemokines as well as reduced phagocytic ability in AM with chronic drinking (26, 43). We found AM from the ethanol group generated a hyper-inflammatory cytokine and chemokine response to LPS. This fits well with our earlier studies on splenic macrophage and blood monocyte responses to LPS (31, 32), as well as other studies on long-term ethanol exposure and myeloid cells (38). In response to respiratory syncytial virus (RSV), AM produced significantly higher levels of IL-6, IL-12, and TGFα than controls. Increased production of IL-6 in response to RSV could indicate broad hyper-inflammation. However, as IL-12 and TGFα levels are not increased in control AM after RSV stimulation, we believe the production of these with alcohol to be indicative of a non-specific and improper response. These observations are in line with increased susceptibility to RSV in individuals with alcohol use disorder (10, 11). To further analyze this response to RSV, we profiled the transcriptional profiles of the AM after infection and found, interestingly, a reduced transcriptional response. Moreover, DEG detected only in the control AM mapped to response to interferon pathways, indicating potential disruptions in antiviral pathway responses with chronic ethanol consumption.

Another critical function of AM is resolution of inflammation and tissue repair to avoid complicating conditions like acute respiratory distress syndrome (ARDS) (51). It has been observed that patients with alcohol use disorders have higher risk of developing ARDS (10, 18) and have weakened wound healing capacities (52). RNA-Seq of AM revealed decreased expression of genes mapping to wound healing processes, such as *CLEC1B*, in ethanol AM. Moreover, reduced chromatin accessibility was noted in promoters that regulate genes important for endothelium development and cell junction assembly with ethanol.

The scRNA-Seq revealed significant heterogeneity within tissue resident and monocyte derived macrophage populations. Recent studies on human lung macrophages have also shown significant macrophage diversity with disease states (30). Clusters that were abundant with ethanol consumption exhibited a transcriptional profile consistent with heightened activation and inflammation. Trajectory analysis further showed an accelerated differentiation of monocytes to macrophages with ethanol. Additional studies are needed to confirm determine whether these clusters represent independent lineages or different activation states associated with ethanol drinking Activation and differentiation in the heterogeneous macrophage populations are controlled by complex epigenetic mechanisms (53). It is possible that the heightened activation state as well as differentiation trajectory with ethanol can be attributed to a process akin to innate training where environmental factors lead to epigenetic changes that have long-lasting functional consequences (54).

Functionally, we report reduced phagocytosis of *S. aureus* by AM from the ethanol AM. Reduction in phagocytosis with long term and acute ethanol exposure has been previously observed in cell culture and rodent models (43, 55). It has been shown in culture that one mechanism of impaired phagocytic function of AM with ethanol is oxidative stress induced by increased NADPH oxidase (56-58). While studies have reported link between ethanol and its metabolites and changes in global methylation state, histone modifications, and ROS production, studies in *in vivo* settings have been limited (59). Data presented in this study show increased levels of intracellular reactive oxygen species (ROS) in AM with ethanol. Ethanol metabolism is a cause of oxidative stress and is directly involved in the production of ROS (60). As ROS serve as inflammasome activating signals and induce inflammation (61), this could contribute to increased production of cytokines and chemokines in response to stimulation. Since the mitochondrial respiratory chain complex I is one of the major contributors of cellular ROS (62), we measured mitochondrial content in AM and found increased mitochondria with ethanol. Future studies would be needed to determine the bioenergetics of these mitochondria and whether they are contributing directly to increased ROS levels and further inflammation in AM (63).

This study provides a comprehensive examination of AM in the context of chronic alcohol drinking in macaques. Our findings indicate increased baseline activation and inflammation signatures epigenetically and transcriptionally in AM that could contribute to increased non-specific inflammatory response to pathogens and compromised phagocytic ability. Potential new targets identified here include increased mitochondrial content, epigenetic alterations, and increased cathepsins in AM with ethanol drinking. These altered AM states could contribute to the increased susceptibility of patients with alcohol use disorders to respiratory infections.

## METHODS AND MATERIALS

### Animal studies and sample collection

These studies used blood and bronchoalveolar lavage (BAL) samples from 9 female and 8 male rhesus macaques (average age 5.68 yrs), with 7 animals serving as controls and 10 classified as chronic heavy drinkers based on over 12 months of daily ethanol self-administration. These samples were obtained through the Monkey Alcohol Tissue Research Resource (https://gleek.ecs.baylor.edu/; Cohorts 6 and 7a). Details about this non-human primate model of voluntary ethanol self-administration have been described (64-66). These cohorts of animals were described in three previous studies of innate immune system response to alcohol (31, 32, 36). BAL cells were obtained after 12 months of open access (22 hr/day alcohol availability) and centrifuged, pelleted and cryopreserved until they could be analyzed as a batch. The average daily ethanol intake for each animal is outlined in **Supp. Table 1**.

### Flow cytometry analysis

1-2×10^6^ BAL cells were stained with the following surface antibodies (2 panels) against: CD206 (BD, 19.2), CD169 (Biolegend, 7-239), HLA-DR (Biolegend, L243), CD14 (Biolegend, M5E2), CD11c (Biolegend, 3.9), CD40 (Biolegend, 5C3), CD163 (Biolegend, GHI/61), CD86 (Biolegend, IT2.2), CX3CR1 (Biolegend, 2A9-1), CCR7 (Biolegend, GO43H7), and CCR5 (Biolegend, J418F1). Samples were acquired with an Attune NxT Flow Cytometer (ThermoFisher Scientific, Waltham, MA) and analyzed using FlowJo software (Ashland, OR).

### Monocyte/Macrophage Stimulation Assays

6.5×10^4^ FACS sorted CD206+ cells from the BAL were cultured in RPMI supplemented with 10% FBS with or without 100 ng/mL LPS or respiratory syncytial virus (RSV) at an MOI of 5 for 16 hours, in 96-well tissue culture plates at 37C in a 5% CO_2_ environment. Plates were spun down: supernatants were used to measure production of immune mediators and cell pellets were resuspended in Qiazol (Qiagen, Valencia CA) for RNA extraction. Both cells and supernatants were stored at −80C until they could be processed as a batch.

### Luminex Assay

Supernatants from AM stimulated with LPS or RSV were measured the ProcartaPlex 31-plex panel measuring levels of cytokines (IFNα, IFNβ, IL-1β, IL-10, IL-12p70, IL-15, IL-17A, IL-1RA, IL-2, IL-4, IL-5, IL-6, IL-7, MIF, and TNFα), chemokines (BLC(CXCL13), Eotaxin (CCL11), I-TAC(CXCL11), IL-8(CXCL8), IP-10(CXCL10), MCP-1(CCL2), MIG(CXCL9), MIP-1a(CCL3), MIP-1b(CCL4)), growth factors (BDNF, G-CSF, GM-CSF, PDGF-BB, VEGF-A) and other factors (CD40L, Granzyme B) (Invitrogen, Carlsbad, CA). Differences in induction of proteins post stimulation were tested using both unpaired (Control-EtOH; Welch’s correction) and paired (NS-Stim) t-tests. Dose-dependent responses were modeled based on g/kg/day ethanol consumed and tested for linear fit using regression analysis in Prism (GraphPad, San Diego CA).

### RNA isolation and library preparation

Total RNA was isolated from purified AM using the mRNeasy kit (Qiagen, Valencia CA) following manufacturer instructions and quality assessed using Agilent 2100 Bioanalyzer. Libraries from PBMC RNA were generated using the TruSeq Stranded RNA LT kit (Illumina, San Diego, CA, USA). Libraries from purified CD14+ monocytes RNA were generated using the NEBnext Ultra II Directional RNA Library Prep Kit for Illumina (NEB, Ipswitch, MA, USA), which allows for lower input concentrations of RNA (10ng). For both library prep kits, rRNA depleted RNA was fragmented, converted to double-stranded cDNA and ligated to adapters. The roughly 300bp-long fragments were then amplified by PCR and selected by size exclusion. Libraries were multiplexed and following quality control for size, quality, and concentrations, were sequenced to an average depth of 20 million 100bp reads on the NextSeq platform.

### Bulk RNA-Seq data analysis

RNA-Seq reads were quality checked using FastQC (https://www.bioinformatics.babraham.ac.uk/projects/fastqc/), adapter and quality trimmed using TrimGalore(https://www.bioinformatics.babraham.ac.uk/projects/trim_galore/), retaining reads at least 35bp long. Reads were aligned to *Macaca mulatta* genome (Mmul_8.0.1) based on annotations available on ENSEMBL (Mmul_8.0.1.92) using TopHat (67) internally running Bowtie2 (68). Aligned reads were counted gene-wise using GenomicRanges (69), counting reads in a strand-specific manner. Genes with low read counts (average <5) and non-protein coding genes were filtered out before differential gene expression analyses. Read counts were normalized using RPKM method for generation of PCA and heatmaps. Raw counts were used to test for differentially expressed genes (DEG) using edgeR (70), defining DEG as ones with at least two-fold up or down regulation and an FDR controlled at 5%. Functional enrichment of gene expression changes in resting and LPS-stimulated cells was performed using Metascape (71). Networks of functional enrichment terms were generated using Metascape and visualized in Cytoscape (72). Transcription factors that regulate expression of DEG were predicted using the ChEA3 (39) tool using ENSEML ChIP database.

### 10X 3’ scRNA-Seq

Freshly thawed BAL from control (n=3) and EtOH (n=3) animals were stained with anti-CD14 antibody and sorted for live monocytes/macrophages using FSC/SSC parameters and CD14+ cells on a BD FACSAria Fusion. Sorted AM/monocytes were pooled and resuspended at a concentration of 1,200 cells/ul and loaded into the 10X Chromium gem aiming for an estimated 10,000 cells per sample. cDNA amplification and library preparation (10X v3.1 chemistry) were performed according to manufacturer protocol and sequenced on a NovaSeq S4 (Illumina) to a depth of >30,000 reads/cell.

### scRNA-Seq data analysis

Sequencing reads were aligned to the Mmul_8.0.1 reference genome using cellranger v3.1 (14) (10X Genomics). Quality control steps were performed prior to downstream analysis with *Seurat (73)*, filtering out cells with fewer than 200 unique features and cells with greater than 20% mitochondrial content. Control and EtOH datasets were integrated in *Seurat* using the *IntegrateData* function. Data normalization and variance stabilization were performed, correcting for differential effects of mitochondrial and cell cycle gene expression levels. Clustering was performed using the first 20 principal components. Small clusters with an over-representation of B and T cell gene expression were removed for downstream analysis. Clusters were visualized using uniform manifold approximation and projection (UMAP) and further characterized into distinct AM/monocyte subsets using the *FindMarkers* function (**Supp. Table 3**).

### Blood monocyte/macrophage integration

Seurat objects from BAL AM/monocytes were integrated with blood monocyte data from the same animals (31) using *Harmony* (74) in order to determine the level of blood monocyte infiltration into the alveolar space. Cells from the BAL that clustered more closely with blood monocytes were identified and that information was projected back onto the original UMAP.

### Pseudo-temporal analysis

Pseudotime trajectory of the AM/monocytes was reconstructed using Slingshot (75). The UMAP dimensional reduction performed in Seurat was used as the input for Slingshot. For calculation of the lineages and pseudotime, the blood monocyte cluster was selected as the start. Temporally expressed genes were identified by ranking all genes by their variance across pseudotime and then further fit using GAM with pseudotime as an independent variable.

### Differential expression analyses

Differential expression analysis (EtOH relative to Control) was performed using MAST under default settings in *Seurat*. Only statistically significant genes (Fold change cutoff ≥ 1.2; adjusted p-value≤ 0.05) were included in downstream analysis.

### Module Scoring and functional enrichment

For gene scoring analysis, we compared gene signatures and pathways from KEGG (https://www.genome.jp/kegg/pathway.html) (**Supp. Table 4**) in the AM/monocytes using *Seurat’s AddModuleScore* function. Values for module scores were further exported from *Seurat* and tested for significance in Prism 7. Over representative gene ontologies were identified by enrichment of differential signatures using Metascape. All plots were generated using *ggplot2* and *Seurat*.

### ATAC-Seq library preparation

10^5^ purified CD206+ alveolar macrophages were lysed in lysis buffer (10 mM Tris-HCl (pH 7·4), 10 mM NaCl, 3 mM MgCl_2_, and NP-40 for 10 min on ice to prepare the nuclei. Immediately after lysis, nuclei were spun at 500 *g* for 5 min to remove the supernatant. Nuclei were then incubated with Tn5 transposase and tagmentation buffer at 37C for 30 min. Stop buffer was then added directly into the reaction to end the tagmentation. PCR was performed to amplify the library for 15 cycles using the following PCR conditions: 72C for 3 min; 98C for 30s and thermocycling at 98C for 15 s, 60C for 30s and 72C for 3 min; following by 72C 5 min. Libraries were then purified with AMPure (Beckman Coulter, Brea CA) beads and quantified on the Bioanalyzer (Agilent Technologies, Santa Clara CA). Libraries were multiplexed and sequenced to a depth of 50 million 100bp paired reads on a NextSeq (Illumina).

### ATAC-Seq data analysis

Paired ended reads from sequencing were quality checked using FastQC and trimmed to a quality threshold of 20 and minimum read length 50. Trimmed reads were aligned to the Macaca Mulatta genome (Mmul_8.0.1) using Bowtie2 (-X 2000 -k 1 --very-sensitive --no-discordant --no-mixed). Reads aligning to mitochondrial genome were removed using Samtools and PCR duplicate artifacts were removed using Picard. Samples from each group were concatenated and accessible chromatin peaks were called using Homer’s *findPeaks* function (76) (FDR<0.05) and differential peak analysis was performed using Homer’s *getDifferentialPeaks* function (P < 0.01). Genomic annotation of open chromatin regions in monocytes and differentially accessible regions (DAR) with EtOH was assigned using ChIPSeeker (77). Promoters were defined as −1000bp to +100bp around the transcriptional start site (TSS). Functional enrichment of differentially accessible promoter regions was performed using Metascape.

Due to the lack of available macaque annotation databases, non-promoter regions from the macaque assembly were converted to the human genome (hg38) coordinates using the UCSC liftOver tool. *Cis*-Regulatory roles of these putative enhancer regions were identified using GREAT (http://great.stanford.edu/public/html/). The Washington University Genome Browser was used to visualize pile-ups (https://epigenomegateway.wustl.edu/). Over-representative transcription factor motifs were identified using Homer’s *findMotifs* function with default parameters. A counts matrix was generated for these regions using *featureCounts (78)*, where pooled bam files for each group were normalized to total numbers of mapped reads.

### Phagocytosis Assay

500,000 freshly thawed total BAL cells were resuspended in RP10 media supplemented with 100ng/mL LPS and incubated for 4 hours at 37C with 5% CO_2_. 50uL of pHrodo Red S.aureus BioParticles (Thermo Fisher Scientific, Waltham, MA) were added to the cells and they were incubated for an additional 2 hours in the incubator. The cells were washed and stained with anti-CD206 antibody and acquired with an Attune NxT Flow Cytometer (ThermoFisher Scientific, Waltham, MA) and further analyzed using FlowJo software (Ashland, OR). Staining positive and negative controls were included for surface markers and pHrodo reagent.

### ROS Assay

500,000 freshly thawed total BAL cells were resuspended in RP10 media supplemented with 100ng/mL LPS and incubated for 3 hours at 37C with 5% CO_2_. 250uM CellRox Deep Red Reagent (ThermoFisher, Waltham, MA) was added at the 3-hour mark and left to incubate for an additional 30 min. The cells were washed and stained with anti-CD206 (BD, 19.2), anti-CD169 (Biolegend, 7-239) antibodies and acquired with an Attune NxT Flow Cytometer (ThermoFisher Scientific, Waltham, MA) and further analyzed using FlowJo software (Ashland, OR). Staining positive and negative controls were included for surface markers and CellRox reagent.

### Mitochondria content measurement

500,000 freshly thawed total BAL cells were resuspended in 50 nM MitoTracker (Invitrogen) probe staining solution and incubated for 30 min at 37C with 5% CO_2_. The cells were washed and stained with anti-CD206 (BD, 19.2), anti-CD169 (Biolegend, 7-239) antibodies and acquired with an Attune NxT Flow Cytometer (ThermoFisher Scientific, Waltham, MA) and further analyzed using FlowJo software (Ashland, OR). Staining positive and negative controls were included for surface markers and MitoTracker reagent.

### Statistical Analysis

All statistical analyses were conducted in Prism 7(GraphPad). Data sets were first tested for normality using Shapiro Wilk test. Two group comparisons were carried out using an unpaired t-test with Welch’s correction or a paired t-test. If normal distribution was not achieved, a non-parametric Mann-Whitney test was used. Differences between 4 groups were tested using one-way ANOVA (⟨=0.05) followed by Holm Sidak’s multiple comparisons tests. Error bars for all graphs are defined as ± SEM. Linear regression analysis compared significant shifts in curve over horizontal line, with spearman correlation coefficient or r^2^ reported. Statistical significance of functional enrichment was defined using hypergeometric tests. P-values less than or equal to 0.05 were considered statistically significant. Values between 0.05 and 0.1 are reported as trending patterns.

## Author Contributions

S.A.L., K.A.G., and I.M. conceived and designed the experiments. S.A.L., B.D., and A.J. performed the experiments. S.A.L. and B.D. analyzed the data. S.A.L. and I.M. wrote the paper. All authors have read and approved the final draft of the manuscript.

## Acknowledgements

We are grateful to the members of the Grant laboratory for expert animal care and sample procurement. We thank Dr. Jennifer Atwood for assistance with sorting in the flow cytometry core at the Institute for Immunology, UCI. We thank Dr. Melanie Oakes from UCI Genomics and High-Throughput Facility for assistance with 10X library preparation and sequencing.

## Data availability

The datasets supporting the conclusions of this article are available on NCBI’s Sequence Read Archive (SRA# Pending).

## SUPPLEMENTARY FIGURES

**Supp. Figure 1:**
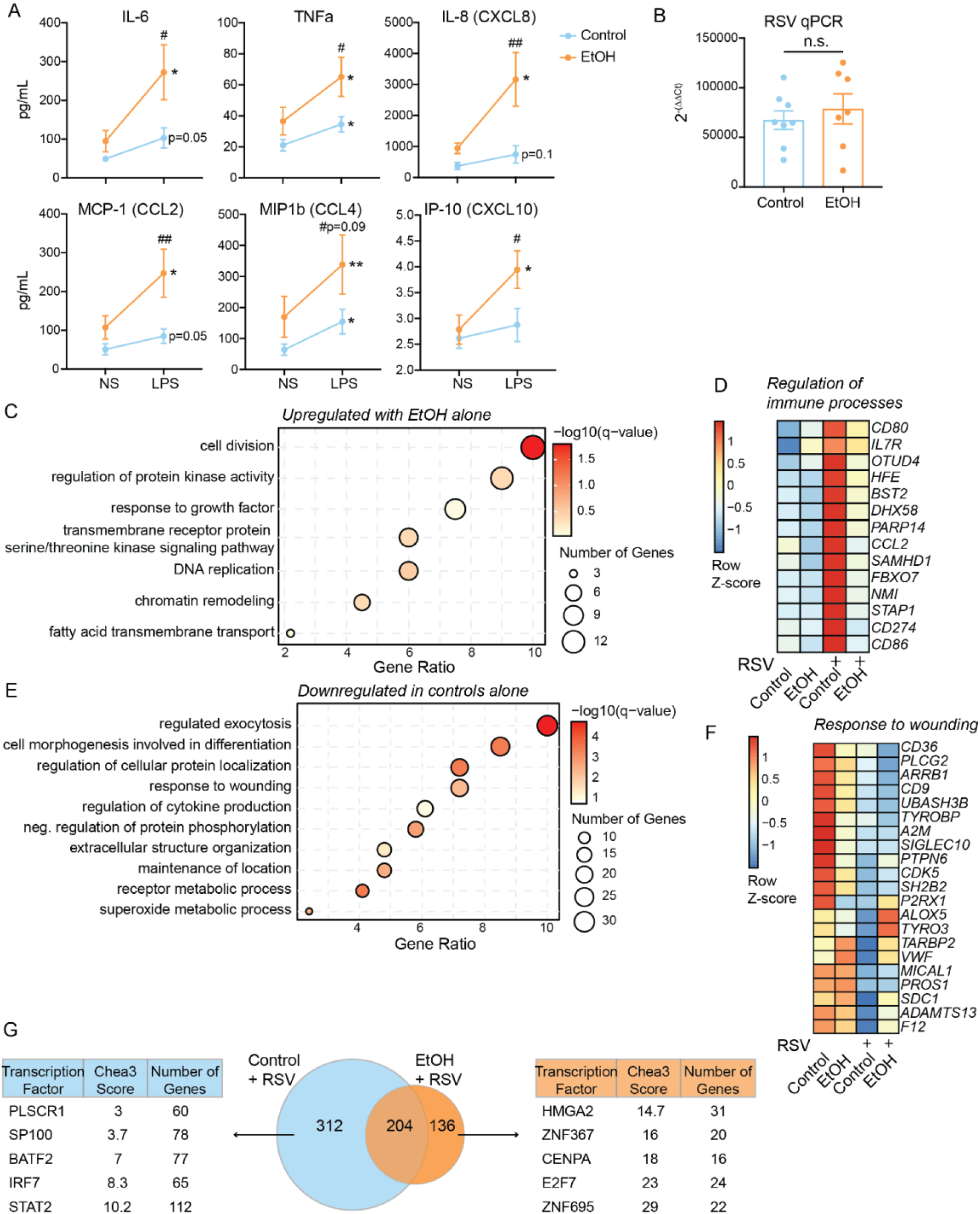
EtOH drinking induces defects in pathogen response in AM. AM were purified and stimulated with LPS for 16 hours. A) Supernatants from LPS stimulation were analyzed by Luminex assay. Line plots representing the measure pg/mL values for the selected analytes. Significance between NS and LPS conditions was tested using a paired t-test and between groups was tested by one-way ANOVA. B) Bar graph of 2^ΔΔCt^ values from qPCR for RSV transcripts after infection. C) Bubble plot showing GO Biological Process enrichment of upregulated DEG in EtOH group alone with RSV stimulation. The size of the bubble represents the number of genes associated with that term, the color represents -log_10_ q-value, and the X-axis is the ratio of genes mapping to that term to total genes. D) Heatmap representing the averaged expression of DEG from *Negative regulation of immune response* term where the scale is Row Z-score representing low (blue) and high (red) expression. E) Bubble plot showing GO Biological Process enrichment of downregulated DEG in control group alone with RSV stimulation. The size of the bubble represents the number of genes associated with that term, the color represents - log_10_ q-value, and the X-axis is the ratio of genes mapping to that term to total genes. F) Heatmap representing the averaged expression of DEG from *Response to wounding* term where the scale is Row Z-score representing low (blue) and high (red) expression. G) Venn diagram comparing upregulated DEG in control and EtOH AM with RSV stimulation. Analysis of the transcription factors regulating the unique DEG was performed using the Chea3 web browser. *=p<0.05, **=p<0.01, ***=p<0.001, ****=p<0.0001. Where indicated # is significance between control and EtOH groups.

**Supp. Figure 2:**
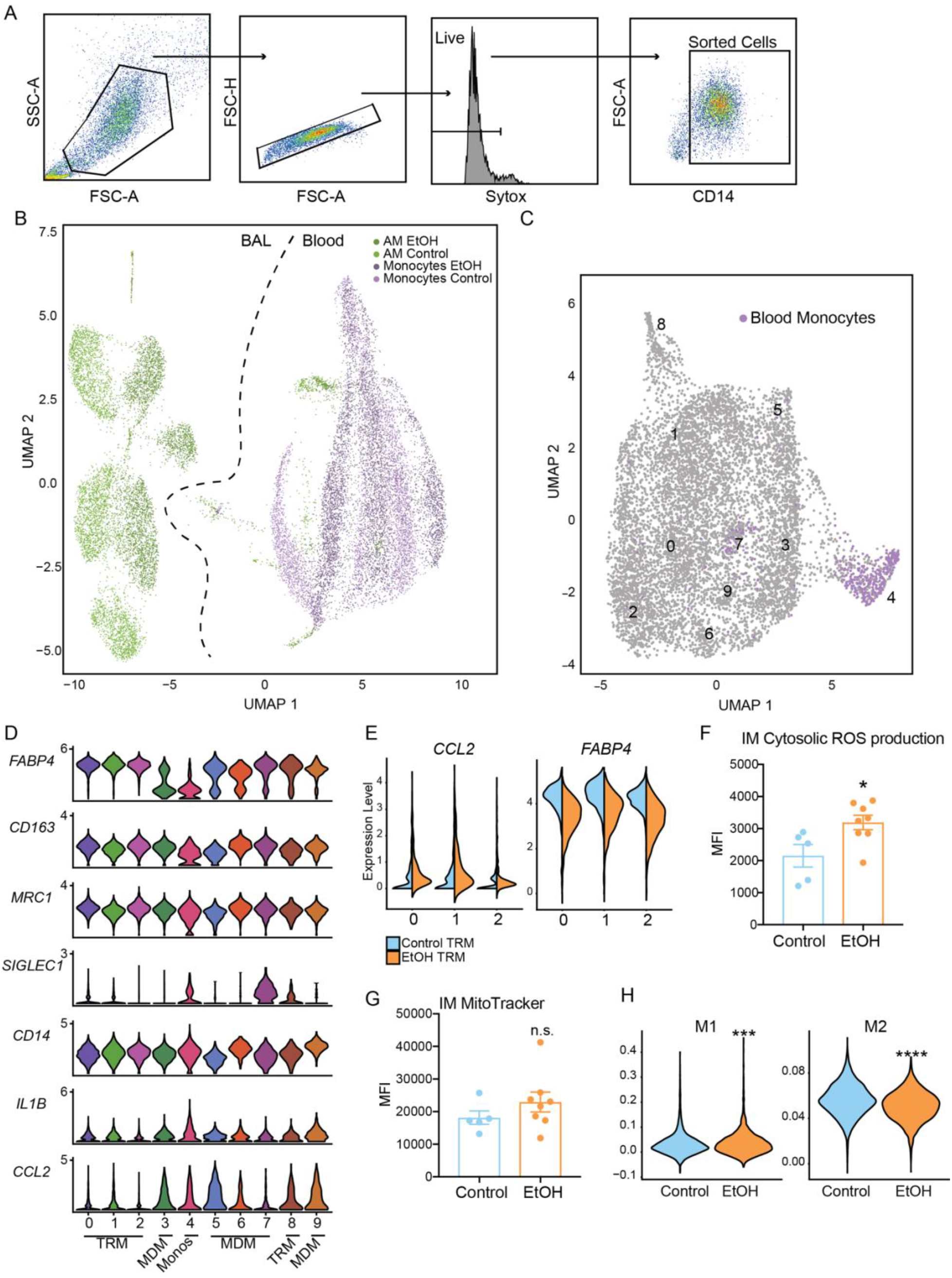
scRNA-Seq and functional implications of EtOH drinking in AM. A) Gating strategy for cell sorting for the 10X scRNA-Seq experiment. B) UMAP of integrated blood (31) and BAL analysis for identification of infiltrating monocytes in the BAL. C) Identified infiltrating blood monocytes highlighted in the original UMAP from Figure 5A. D) Violin plots of markers associated with tissue resident (*FABP4, CD163, MRC1, SIGLEC1*) and infiltrating (*CD14, IL1B, CCL2*) cells. E) Split violin plots of DEG detected between EtOH and control in TRM clusters. F) Bar plot showing median fluorescence intensity (MFI) of intracellular oxidative stress stained by CellROX Green Reagent in IM. G) Bar plot showing median fluorescence intensity (MFI) of intracellular mitochondria stained with MitoTracker Red in IM. H) Violin plots representing M1 and M2 module scoring of total cells from each group. Significance was calculated using Mann-Whitney test. *=p<0.05, **=p<0.01, ***=p<0.001, ****=p<0.0001.

## SUPPLEMENTARY TABLES

**Supp. Table 1**: Animals and EtOH g/kg values

**Supp. Table 2**: Immune mediator production by AM following LPS or RSV stimulation

**Supp. Table 3**: Genes associated with each cluster

**Supp. Table 4**: Module Scoring genes

