## Supplementary material for "Chronic ethanol drinking in non-human primates induces inflammatory cathepsin gene expression in alveolar macrophages accompanied by functional defects": Supp Table 1

| **MATRR ID** | **Sex** | **Mean daily ethanol intake (g/kg/day)** | **Standard Deviation of mean daily intake** | **Blood**  **Ethanol**  **Content**  **(Avg mg%)** | **Drinking status** |
| --- | --- | --- | --- | --- | --- |
| 10068 | F | 0 | - | 0 | Control |
| 10071 | F | 0 | - | 0 | Control |
| 10076 | F | 0 | - | 0 | Control |
| 10186 | F | 0 | - | 0 | Control |
| 10187 | F | 0 | - | 0 | Control |
| 10093 | M | 0 | - | 0 | Control |
| 10094 | M | 0 | - | 0 | Control |
| 10095 | M | 0 | - | 0 | Control |
| 10096 | M | 0 | - | 0 | Control |
| 10080 | F | 3.3 | 1.0 | 41 | EtOH |
| 10070 | F | 3.9 | 1.0 | 46 | EtOH |
| 10081 | F | 4.0 | 1.1 | 57 | EtOH |
| 10079 | F | 4.0 | 1.3 | 65 | EtOH |
| 10078 | F | 5.0 | 1.4 | 81 | EtOH |
| 10069 | F | 5.2 | 1.2 | 103 | EtOH |
| 10088 | M | 2.9 | 1.3 | 74 | EtOH |
| 10091 | M | 3.2 | 0.9 | 79 | EtOH |
| 10097 | M | 3.0 | 0.6 | 79 | EtOH |
| 10098 | M | 3.3 | 1.0 | 98 | EtOH |

**Table 1: Summary of samples used in this study.** Mean daily ethanol (EtOH) intake reflects the average dose consumed during the period of 12-month self-administration period. Blood for ethanol concentration was taken at 7 hrs into the 22 hr daily session every 5-7 days throughout the 12 consecutive months of alcohol access and analyzed by gas chromatography.
